# Role of Cockayne Syndrome B (CSB) Protein in Genome Maintenance in Human Cells under Oxidative Stress

**DOI:** 10.1101/2025.01.02.631021

**Authors:** Grace Kah Mun Low, Gavin Yong-Quan Ng, Dimphy Zeegers, Aloysius Ting, Kalpana Gopalakrishnan, Aik Kia Khaw, Manikandan Jayapal, M Prakash Hande

**Affiliations:** Department of Physiology, Yong Loo Lin School of Medicine, National University of Singapore, Singapore

**Keywords:** Cockayne Syndrome B (CSB), Genome Stability, Oxidative Stress, Transcription Coupled Repair (TCR), Reactive Oxygen Species (ROS), Nucleotide Excision Repair (NER), Telomere Dynamics, Ageing

## Abstract

Cockayne Syndrome (CS), a disorder marked by premature ageing and neurodevelopmental abnormalities, is primarily caused by mutations in the CSB protein, a critical component of the transcription-coupled repair pathway of nucleotide excision repair. This study explores the role of CSB in managing oxidative DNA damage and maintaining telomere integrity under oxidative stress conditions. We subjected CSB-deficient human fibroblasts (CS-B) and control fibroblasts to acute and chronic oxidative stress through hydrogen peroxide (H₂O₂) treatment and elevated oxygen levels. Our findings reveal that CS-B fibroblasts exhibit a distinct resistance to acute oxidative stress, as evidenced by their sustained viability and minimal cell cycle arrest compared to control fibroblasts. However, chronic oxidative conditions led to accelerated senescence in CS-B fibroblasts, demonstrated by increased telomere attrition rates, senescent morphology, and upregulated senescence-associated β-galactosidase activity. Further, gene expression analysis post-H₂O₂ exposure identified the downregulation of key DNA repair and cell cycle genes in CS-B fibroblasts, suggesting a compromised ability to respond to oxidative DNA damage. These observations underscore the multifaceted role of CSB in genomic maintenance and highlight its potential involvement in the pathology of CS through impaired response to oxidative stress and telomere instability. The study contributes to a deeper understanding of the cellular mechanisms that underlie CS symptoms and may inform potential therapeutic strategies targeting oxidative damage repair systems.

## 1.0 Introduction

Cockayne Syndrome (CS) typically results from mutations in the Cockayne syndrome protein A (*CSA/ERCC 8*) or Cockayne syndrome protein B (*CSB/ERCC 6*) genes and is thus a transcription-coupled repair (TCR)-nucleotide excision repair (NER) specific disorder [1]. Approximately 80 % of CS cases are caused by a defect in CSB [2]. The CSB/ERCC6 protein is 168 kDa in size and is involved in the TCR sub-pathway of the NER [3, 4] where DNA lesions from transcribed strands of active genes are preferentially excised to restore UV-inhibited transcription rapidly. CSB is suggested to recognise and displace stalled RNA polymerase II and recruit other NER proteins for subsequent steps in the repair process [4–6]. CSB has also been implicated in the base excision repair (BER) of oxidative lesions such as 8-oxoguanine and 8-oxoadenine via interactions with DNA glycosylases in the nucleus and mitochondria [7–9].

It has been suggested that CSA and CSB both play roles in basal transcription or other forms of repair, and different mutations result in subtle impairment in transcription, leading to developmental abnormalities [10–12]. During normal developmental processes, endogenous oxidative stress-induced DNA lesions coupled with inefficient transcription could impair physical and neurological development, leading to dwarfism and demyelination [11]. In support of this, CS cells exhibited a defect in TCR of oxidative DNA lesions and are slightly more sensitive to oxidative damage-inducing ionising radiation than normal cells [13]. Consistent with this, the CSB protein has been implicated in post-oxidative stress transcriptional response. Cells defective in CSB failed to induce ribosomal protein expression and exhibited down-regulation of several proteins related to stress response, transcription, translation, signal transduction and cell cycle progression following H2O2 exposure [14].

Increasing studies demonstrate that defects in DNA repair factors such as the RecQ helicases, WRN and BLM,[15–18] and the NER factor XPF/ERCC1 [19, 20] are linked to telomere attrition associated with premature senescence, telomere-related chromosomal anomalies and progeroid features [21–24]CSB-deficient mice, like CS patients, present with premature ageing features associated with increased sensitivity to oxidative stress. Since oxidative stress accelerates telomere shortening, we hypothesise that CSB is involved not only in DNA repair, oxidative stress management, and UV-inhibited transcription but also in telomere homeostasis.

As such, in this study, we sought to determine the involvement of CSB in oxidative DNA damage-repair and telomere dynamics by treating primary fibroblasts from a CS-B patient, hereafter known as CS-B fibroblasts, with H2O2 or increased ambient O2 partial pressure of 40% (40% O2). We performed assays related to survival, genome stability and growth kinetics. Our study has presented us with a viewpoint on how CSB protein is involved in genomic maintenance during oxidative stress in human cells.

## 2.0 Materials & Methods

### 2.1 Cell Culture and Treatment Conditions

Primary human diploid fibroblasts from normal individuals (GM03651E and IMR-90 termed as *Normal* and I90 for this study), and from individuals suffering from Cockayne Syndrome Type B (CS-B GM00739) were purchased from Coriell Cell Repository and cultured in Minimal Essential Medium (MEM; Gibco, U.S.A.) supplemented with 15% fetal bovine serum ( Hyclone, U.S.A.), 100 U/mL penicillin/streptomycin, 1% vitamins, and 2% essential and 1% non-essential amino acid. All cells were grown in a humidified 5% CO2 incubator at 37°C and maintained in a log phase.

A fresh working solution of 100 mM H_2_O_2_ (Kanto Chemical Co. Inc., Japan) was prepared in plain media each time. For dose-response studies, exponentially growing cells were exposed to between 20 µM and 100 µM of H2O2 for 2 hours, following which the medium was replaced with fresh medium for a 22-hour recovery period. Such concentrations have been shown to be of low to no cytotoxicity with some genotoxicity.

For long-term studies, approximately 2 × 10^5^ cells were seeded in T75 (*Nunc*) tissue culture flasks. Cells were subjected to a 30-day low-dose chronic treatment. Cells were either treated with 10 μM H_2_O_2_ every 2 days with media changed prior to each treatment, or with 20 µM H_2_O_2_ every 2 days with media changed prior to each treatment or were maintained in a 40% O_2_ incubator with media changed every 2 days. A parallel set of cells had a media change every 2 days when grown in standard culture conditions.

### 2.2 Crystal Violet Assay for Cell Viability

After treatment, the attached cells were gently washed in 1x PBS. Crystal Violet solution (0.75% Crystal Violet (Sigma) in 50% ethanol (Merck, Germany), distilled H_2_O (dH2O), 1.75% v/v formaldehyde (Merck), and 0.25% w/v sodium chloride (NaCl; Lab-Scan Ireland)) was carefully added to the wells and incubated at room temperature, followed by a wash with 1x PBS.. The wells were then dried at 37°C. To lyse the cells and to solubilize the dye, a solution of 1% sodium dodecyl sulphate (SDS; NUMI supplies) in 1x PBS was added. Absorbance was measured at 595 nm with µQuant Microplate Spectrophotometer (Bio-Tek Instruments Inc., U.S.A). Data is represented as the percentage of absorbance of the sample compared to the untreated control.

### 2.3 Cytokinesis Blocked Micronucleus(CBMN) analysis

After the treatment, cells were incubated with 4.0 µg/mL cytochalasin B (*Sigma*) in fresh medium for 22 hours. The protocol used is adapted from Low et al [25, 26].. One thousand acridine orange stained binucleated cells (BN) with/without the presence of micronuclei (MN) were scored under the Axioplan 2 imaging fluorescence microscope (*Carl Zeiss, Germany*) with appropriate triple band filter.

### 2.4 Alkaline single cellgelelectrophoresis (SCGE/Comet) assay

Following treatment, one set of cells was allowed to undergo a recovery period in fresh media without drug (22 hours for H₂O₂, and 24 hours for arsenite) whilst the other set was harvested immediately. Details of the protocol was explained in our earlier publications [25, 26]. One hundred randomly selected cells per sample were examined using an Axioplan 2 imaging fluorescence microscope and analysed with Comet Imager Software (Metasystems, Germany). The extent of DNA damage was quantified by measuring comet tail moments, which represent the fraction of DNA in the comet tail.

### 2.5 Populationdoubling (PD)

Approximately 2 × 10^5^ cells were seeded in T75 (*Nunc*) tissue culture flasks. Cells were subjected to a 30-day low-dose chronic treatment. Cells were either treated with 10µM H_2_O_2_ every 2 days with media changed prior to each treatment, with 20 µM H_2_O_2_ every 2 days with media changed before each treatment or were maintained in a 40% O_2_ incubator with media changed every 2 days. A parallel set of cells had a media change every 2 days when grown in standard culture conditions. Cells were harvested on days 6, 12, 18 or 30 and counted using a haemocytometer. A fresh tissue culture flask was then reseeded with 2 × 10^5^ cells or all the cells if cell numbers were less than 2 × 10^5^

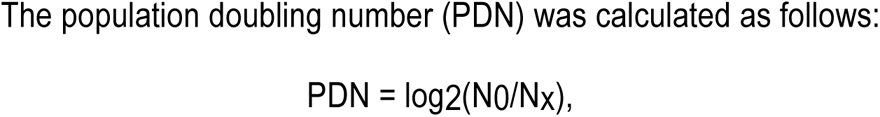

where N0 = number of cells at harvest and Nx= number of cells seeded.

### 2.6 Morphology by lightmicroscopy and cell size

Before media changes, cells were observed under a light microscope at 40× and 100× magnifications. The morphology of the cells was photographed using an Olympus C-7070 WZ digital camera (*Japan*). Cells were shown to increase in size during senescence [27] On days of harvest during the PD study, trypsinised cells were observed under a light microscope and measured for cell diameter using a micrometre eyepiece.

### 2.7 Senescence Associated β-Galactosidase(SA-βgal) Staining

On days 1, 6, 12, 18, and 24 of the PD study, cells were seeded in 6-well plates (Nunc) and assessed for the senescence marker SA-β-gal using the Senescence β-Galactosidase Staining Kit (Cell Signaling Technology) following the manufacturer’s instructions. Attached cells were washed twice with 1× PBS, fixed with a solution of 2% formaldehyde and 0.2% glutaraldehyde in 1× PBS for 15 minutes at room temperature, washed twice with 1× PBS, and incubated overnight at 37°C with 1 mL of staining solution (40 mM citric acid/sodium phosphate, 0.15 M NaCl, 2 mM magnesium chloride) containing 5 mM potassium ferrocyanide, 5 mM potassium ferricyanide, and 1 mg β-gal in N-N-dimethylformamide (Sigma). Cells were observed under a light microscope for a bluish stain indicative of SA-β-gal presence. Photographs were taken at 40×, 100×, and 200× magnifications using an Olympus C-7070 WZ digital camera.

### 2.8 Telomere Length Measurement by Terminal Restriction Fragment (TRF)

DNA was extracted from cells using the DNeasy Tissue Kit (Qiagen, USA) following the manufacturer’s protocol. Telomere restriction fragment (TRF) length was analysed using the Telo-TAGGG Length Assay Kit (Roche Applied Science, USA). Quantitative measurements of the mean TRF length were calculated using the Kodak Gel imaging system and Kodak imaging software. Telomere signals were exposed to X-ray film (Pierce), and their lengths were determined by comparing the signals to a molecular weight marker standard with Kodak Molecular Imaging Software (Carestream Molecular Imaging, USA). The details were followed as reported earlier [28] The decrease in telomere length was expressed as a function of PDN to assess the rate of telomere shortening.

### 2.9 Gene Expression Studies

Total RNA was extracted from the cells using the QIAmp RNA Blood Mini Kit (Qiagen, Germany). The extracted RNA was quantified with a NanoDrop 1000 (Thermo Scientific, USA), and its integrity was checked using a Bio-Analyzer (Agilent Technologies, Inc., USA). For the gene expression study, 750 nanograms of extracted RNA from each sample were used. The TotalPrep RNA Amplification Kit (Ambion Inc., TX, USA) was employed for the cRNA amplification process. The resulting biotinylated amplified RNA was hybridised with HumanRef8 V3.0 Human Whole-Genome Expression BeadChips (Illumina Inc., USA) for 16 hours at 58°C. After incubation, the arrays were washed and stained with Streptavidin-Cy3 (GE Healthcare, Bio-Sciences, UK). The arrays were then scanned using an Illumina Bead Array Reader. The scanned array data were imported and analysed with GeneSpring GX7.3 (Agilent Technologies, USA). Differentially expressed genes (p<0.05, one-way ANOVA) were annotated according to Gene Ontology-Biological ProcessData analysis also involved agglomerative average-linkage hierarchical clustering to determine varied patterns and gene expression levels following treatment with H_2_O_2_ or under 40% Oxygen. Data visualisation tool is constructed using SRplot online [29].

### 2.10 Statistical Analysis

Statistical significance between and among data sets was assessed using two-way ANOVA, using GraphPad Prism. The difference was considered to be statistically significant when *P < 0.05; **P<0.01; and ***P<0.001.

## 3.0 Results

### 3.1 H₂O₂ treatment decreases cell viability of cells, but CSB-deficient fibroblasts show less sensitivity

The cell viability of all three types of fibroblasts decreased in a dose-dependent manner following H₂O₂-treatment (P < 0.05; **Figure 1A**). However, the control fibroblasts, IMR-90 and *Normal*, exhibited lower viabilities than CS-B from concentrations of 60 µM and higher (P < 0.01). Specifically, while the viabilities of IMR-90 and *Normal* fibroblasts were less than 50% at H₂O₂ doses above 60 µM, these concentrations remained sub-lethal to CS-B fibroblasts (**Figure 1A**).

**Figure 1:**
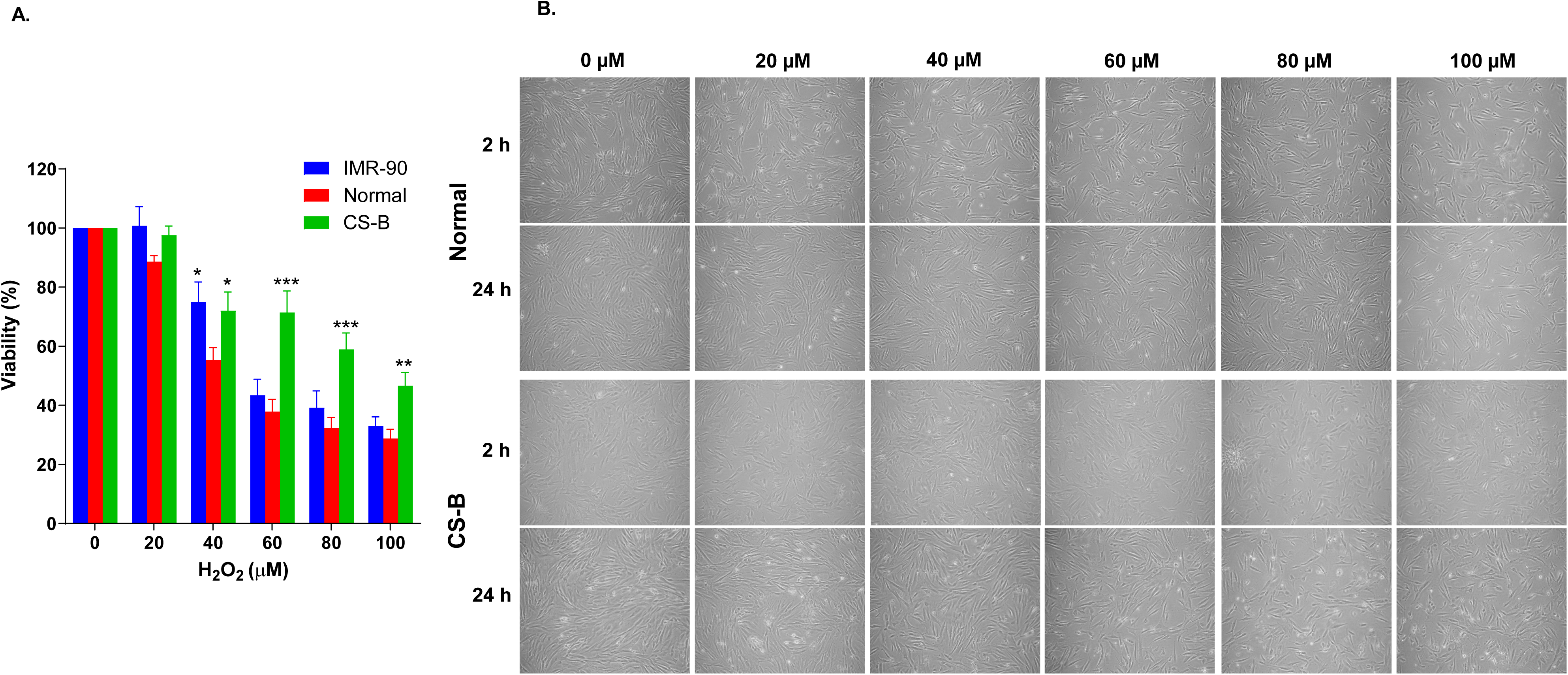
**A:** Cell viability by Crystal Violet assay. Dose-dependent cell death in H₂O₂ -treated fibroblasts. CS-B is significantly less sensitive to cell death compared to control cells at higher concentrations. \**P* < 0.05, \*\**P* < 0.01 compared to normal/control cell types; (^#^*P* < 0.05, ^##^*P* < 0.01; ^###^*P* < 0.001 compared to the untreated counterparts). Two-way ANOVA was used in the statistical analysis. **B:** Morphology of fibroblasts following treatment with H₂O₂ at 40× magnification.

### 3.2 CS-B fibroblasts display minimal morphological changes following H₂O₂ treatment

Morphology of attached fibroblasts was observed two h following H₂O₂ treatment and 22 h after a change of fresh media without H₂O₂ (**Figure 1B**). At the 2-hour timepoint, *Normal* cells exhibited a response to H₂O₂, evident from the observable changes in cell refractivity. However, the morphology of CS-B cells did not change significantly immediately after treatment. Importantly, following 22 h after the fresh change of media treated *Normal* cells were less able to proliferate to confluency compared to untreated counterparts. In contrast, CS-B cells showed signs of cell damage at concentrations of 80 µM and above (**Figure 1B**).

### 3.3 CS-B cells produced significantly more micronuclei than *Normal* cells following H₂O₂ treatment

We next examined the cytokinesis-block micronucleus assay to ascertain if the changes in both cell death and cell cycle profile (data not shown) were associated with DNA damage. Both the frequency of BN with MN and the MN frequency of both *Normal* and CS-B fibroblasts increased significantly with increasing H₂O₂ concentration (P < 0.05; **Table 1** and **Figures 2A & 2B**). H₂O₂ concentrations of 60 µM and higher did not yield enough BN fibroblasts for accurate analysis. Nonetheless, low doses of H₂O₂ (20 and 40 µM) were sufficient to increase significantly both the percentage of BN with MN and the MN frequency of CS-B fibroblasts compared to that of *Normal* fibroblasts (P < 0.05; **Table 1** and **Figures 2A & B**). To note, the occurrence of CS-B BN harbouring two or more MN following 40 µM H₂O₂-treatment was higher than that of its *Normal* counterpart (**Table 1**).

**Figure 2:**
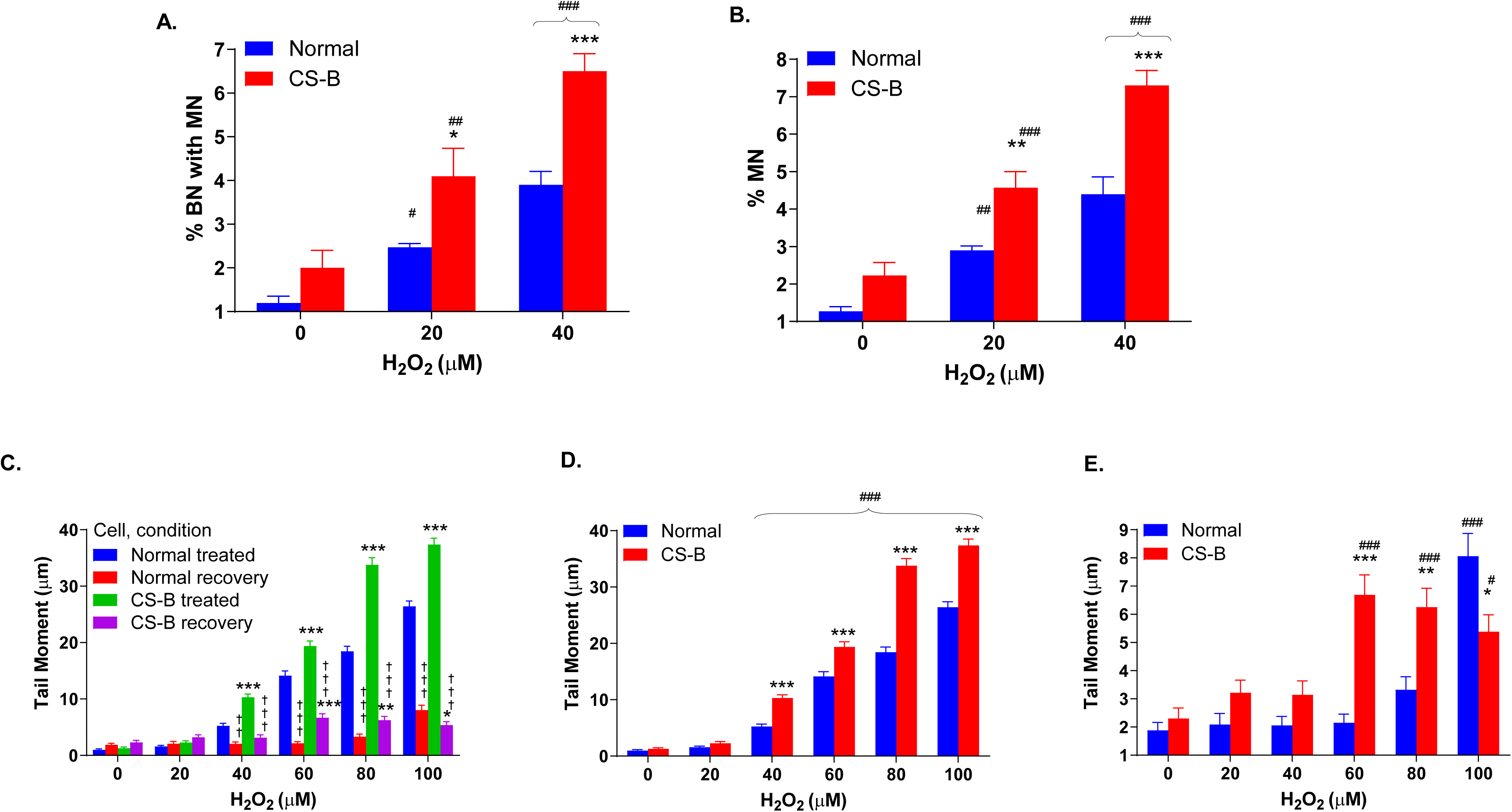
Cytokinesis Blocked Micronucleus (CBMN) analysis **A)** Percent BN with the presence of MN and **B)** Percent MN binucleated fibroblasts scored following H₂O₂ treatment. Average ± S.D. *P < 0.05, **P < 0.01 compared to normal cells; (#P < 0.05, ##P < 0.01; ###P < 0.001 compared to the untreated counterparts). Two-way ANOVA was used in the statistical analysis. **C – E**: Single Cell Gel Electrophoresis of H₂O₂ -treated fibroblasts. **C)** Tail moments immediately following 2 h H₂O₂ treatment and 22 h recovery in fresh medium. Treatment of 40 µM H₂O₂ and above resulted in significantly increased tail moment (P < 0.001) which decreased significantly following recovery (P < 0.001) in both cell types. While there was no significant difference in moments between *Normal* and CS-B cells for both treated and recovering cells at 0 µM and 20 µM H₂O₂ (P > 0.05), CS-B cells exhibited an increase in moments compared to the *Normal* cells for both treated and recovering cells at higher concentrations. **D)** Tail moments immediately following 2 h H₂O₂ treatment. Both cell types exhibit dose-dependent increase in tail moment (P < 0.001). **E)** Tail moments after 22 h recovery in fresh medium. *P < 0.05, **P < 0.01, ***P < 0.001 indicate significantly greater tail moment when comparing CS-B cells to their *Normal* counterparts. ††P < 0.01, †††P < 0.001 indicate significant reduction in the tail moment at recovery from treated. ##P < 0.05, ##P < 0.01, ###P < 0.001 indicate significantly increased tail moments from untreated counterparts. Data is represented as mean ± SE.

**Table 1.** Micronuclei (MN in cytokinesis-blocked binucleated (BN) human fibroblasts following H_2_O_2_ treatment.

|  |  | BN with respective number of MN |  |  |  |  | BN with MN | Total MN | Total MN (% ± SD) |
| --- | --- | --- | --- | --- | --- | --- | --- | --- | --- |
|  | H<sub>2</sub>O<sub>2</sub> (µM) | Total BN scored | 1 MN | 2 MN | 3 MN | 4 MN |  |  |  |
| Normal | 0 | 3000 | 34 | 2 | 0 | 0 | 36 (1.20) | 38 | 1.27 ± 0.2 |
|  | 20 | 3000 | 63 | 9 | 2 | 0 | 74 (2.47) * | 87 | 2.90 ± 0.2** |
|  | 40 | 3000 | 105 | 10 | 1 | 1 | 117 (3.90) *** | 132 | 4.40 ± 0.8*** |
| CS-B | 0 | 3000 | 53 | 7 | 0 | 0 | 60 (2.00) | 67 | 2.23 ± 0.59 |
|  | 20 | 3000 | 113 | 7 | 2 | 1 | 123 (4.10) ** | 137 | 4.57 ± 0.74*** |
|  | 40 | 3000 | 177 | 12 | 6 | 0 | 195 (6.50) *** | 219 | 7.30 ± 0.69*** |

Data is representative of three experiments where 1000 BN were scored at each time. Data is represented as mean ± S.D. \* $P < 0.05$ , where significance is calculated to the cell type's untreated counterpart. \*\* $P < 0.01$ , where significance is calculated to the cell type's untreated counterpart. \*\*\* $P < 0.001$ , where significance is calculated to the cell type's untreated counterpart.

### 3.4 Lack of functional CSB increases DNA damage susceptibility to H₂O₂ and compromises H₂O₂-induced DNA lesion-repair

Next, to study and compare the competence of these cells to recover from H₂O₂-induced DNA damage, we measured the overall genomic integrity of both *Normal* and CS-B fibroblasts using comet tails analysis at two-time points: the first one immediately following the 2-h H₂O₂ treatment and the second one after 22 h of recovery in fresh media without H₂O₂. At the 2-h time point, both *Normal* and CS-B fibroblasts exhibited dose-dependent increases in tail moments at concentrations of 40 µM and higher (P < 0.001; **Figures 2C** & **2D**). Following the 22-h recovery period, both *Normal* and CS-B cells showed significant drops in tail moments compared to their corresponding 2-h counterparts (P < 0.01; **Figure 2C**), suggesting that increasing the concentrations of H₂O₂ induced more DNA damage and that DNA repair took place when cells were allowed to recover. In addition, untreated and 20 µM H₂O₂-treated CS-B fibroblasts did not display any significant differences in tail moments from that of *Normal* counterparts at the 2-h time point (P > 0.05). However, at the same time point, concentrations of 40 µM and above induced greater tail moments in CS-B fibroblasts than in *Normal* counterparts (P < 0.001; **Figure 2D**). Following recovery, although both *Normal* and CS-B fibroblasts demonstrated significantly decreased tail moments compared to their 2-h counterparts (P < 0.01; **Figure 2C**), the recovered tail moments of CS-B did not reduce to that of untreated counterparts from concentrations of 60 µM and up (P <0.05; **Figure 2E**). In addition, at 60 µM and 80 µM H₂O₂, these recovered tail moments were longer than that of *Normal* counterparts (P < 0.001; **Figure 2E**).

### 3.5 Chronic exposure to 10 µM H_2_O_2_ results in senescence and telomere shortening in fibroblasts, whereas CS-B cells exhibit earlier senescent characteristics and increased telomere attrition

We first looked at the population doubling number (PDN) over a period of 6 days to understand the growth rate of the fibroblasts and found that CS-B fibroblasts required a longer time to double compared to control cells (**data not shown)**. We then cultured these cells for 30 days under chronic oxidative stress with a paired control cultured under standard conditions. Consistent with this, untreated CS-B cells displayed slower doubling than *Normal* cells. Treatment with 10 µM H₂O₂ resulted in a reduction of PDN in both cell types, although they continued to show the ability to proliferate (**Figure 3A**). Chronic treatment with 10 µM H₂O₂ accelerated the appearance of senescent morphology of both *Normal* and CS-B cells at days 23 and 22, respectively (**data not shown**).

**Figure 3:**
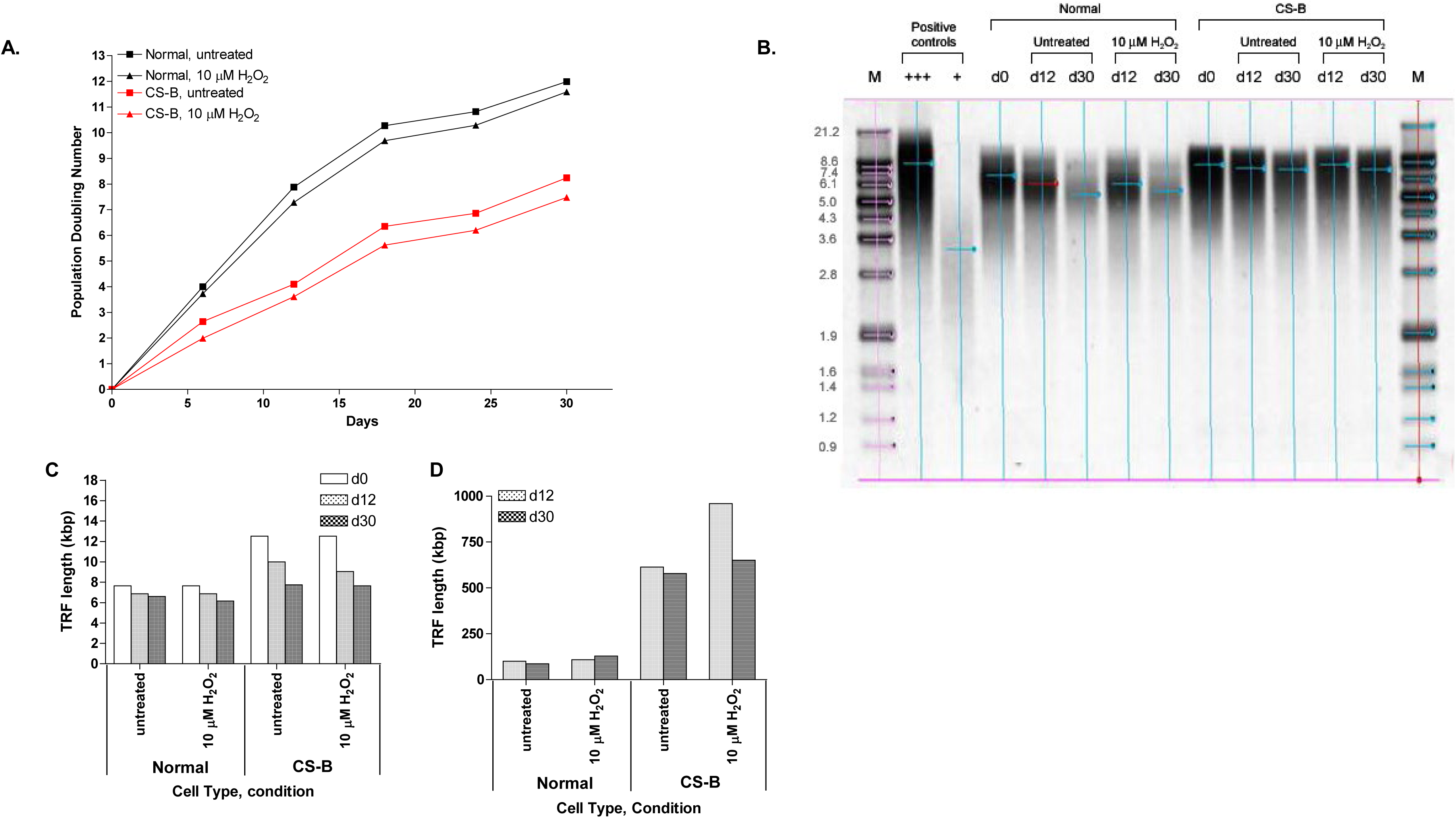
Long-term study of fibroblasts undergoing 30 days of chronic oxidative stress in the form of 10 µM H₂O₂ treatment every 48 h. **A)** Growth kinetics of fibroblasts as depicted by population doubling number (PDN) under chronic oxidative stress of 30 days. PDN is determined by cell counting at intervals of 6 days. Cells treated with H₂O₂ show reduced PDN compared to that of untreated counterparts. CS-B cells display slower growth kinetics compared to that of normal counterparts. **B)** Southern blotting of TRF obtained from a genomic DNA digest using Hinf I and Rsa I restriction enzymes. TRF length at start of treatment and following 12 and 30 days respectively under chronic oxidative stress. TRF length decreases with time for *Normal* and CS-B cells in under all conditions following 30 days. **C)** Rate of TRF length attrition following 12 and 30 days of chronic oxidative stress, respectively. TRF attrition rate is derived by dividing the decrease in TRF length (B by PDN (3A).

Next, total DNA was extracted from cells at the beginning (day 0), middle (day 12) and end (day 30) of the treatment and subjected to terminal restriction fragment (TRF) length analysis, where non-telomeric and non-subtelomeric DNA were digested using Hinf I and Rsa I restriction enzymes. An image of the southern blot of the TRFs from both *Normal* and CS-B fibroblasts is portrayed (**Figure 3B**) Over the 30-day period, both untreated and treated *Normal* and CS-B cells showed a progressive decrease in TRF (**Figure 3C**) Interestingly, the slope of TRF length decrease, indicative of accelerated telomere attrition, is steeper in CS-B samples than that of *Normal* fibroblasts (**Figure 3C**). This is made clear when we looked at the decrease in TRF length at day 12 and day 30 from day 0 of the treatment. Specifically, regardless of exposure to H₂O₂ and of the day of analysis, CS-B fibroblasts showed distinctly increased loss of TRF length compared to that of *Normal* fibroblasts (**Figure 3D)**. Also to note is that treated *Normal* fibroblasts compared to their untreated counterparts exhibited an increased reduction in TRF length after 30 days while treated CS-B fibroblasts showed a more obvious reduction only at 12 days but not 30 days compared to their untreated counterparts.

As telomeres are lost with each cell division, we took the PDN into account to yield TRF attrition rate **Figures 3C and D**) to see if CS-B cells were more susceptible to losing telomeres with each cell doubling. Chronic exposure to such a low dose of H₂O₂ resulted only in a marginal increase in rate of TRF loss in *Normal* fibroblasts. CS-B fibroblasts however showed a distinct increased rate of TRF length attrition following chronic exposure to 10 µM H₂O₂ for the first 12 days of treatment. Notably, CS-B cells had longer telomeres than *Normal* fibroblasts at the beginning of the treatment **(Figure 3D**) and spontaneously lost more telomeres as depicted by the increased rate of TRF attrition in untreated CS-B cells compared to that of *Normal* cells (**Figure 3D**).

### 3.6 Chronic exposure to 20 µM H₂O₂ and 40 % O_2_ results in senescence and telomere shortening in fibroblasts, whereas CS-B cells exhibit earlier senescent characteristics and increased telomere attrition

Cells were exposed chronically to 20 µM H₂O₂ and 40 % O_2_ for 30 days, where they were harvested and counted when untreated cells reached 90 % confluence. IMR-90 fibroblasts showed a much higher proliferation capacity than *Normal* and CS-B fibroblasts regardless of treatment condition as shown by the higher PDN at each time point compared to the other cell types (**Figure 4A**). Importantly, chronic oxidative stress in the form of 20 µM H₂O₂ and 40 % O_2_ both considerably decreased the doubling capacity of all three cell types (**Figure 4A**). Untreated and treated *Normal* and CS-B fibroblasts showed similar growth kinetics, whereas treated fibroblasts hardly doubled over the 30 days of treatment (**Figure 4A**).

**Figure 4:**
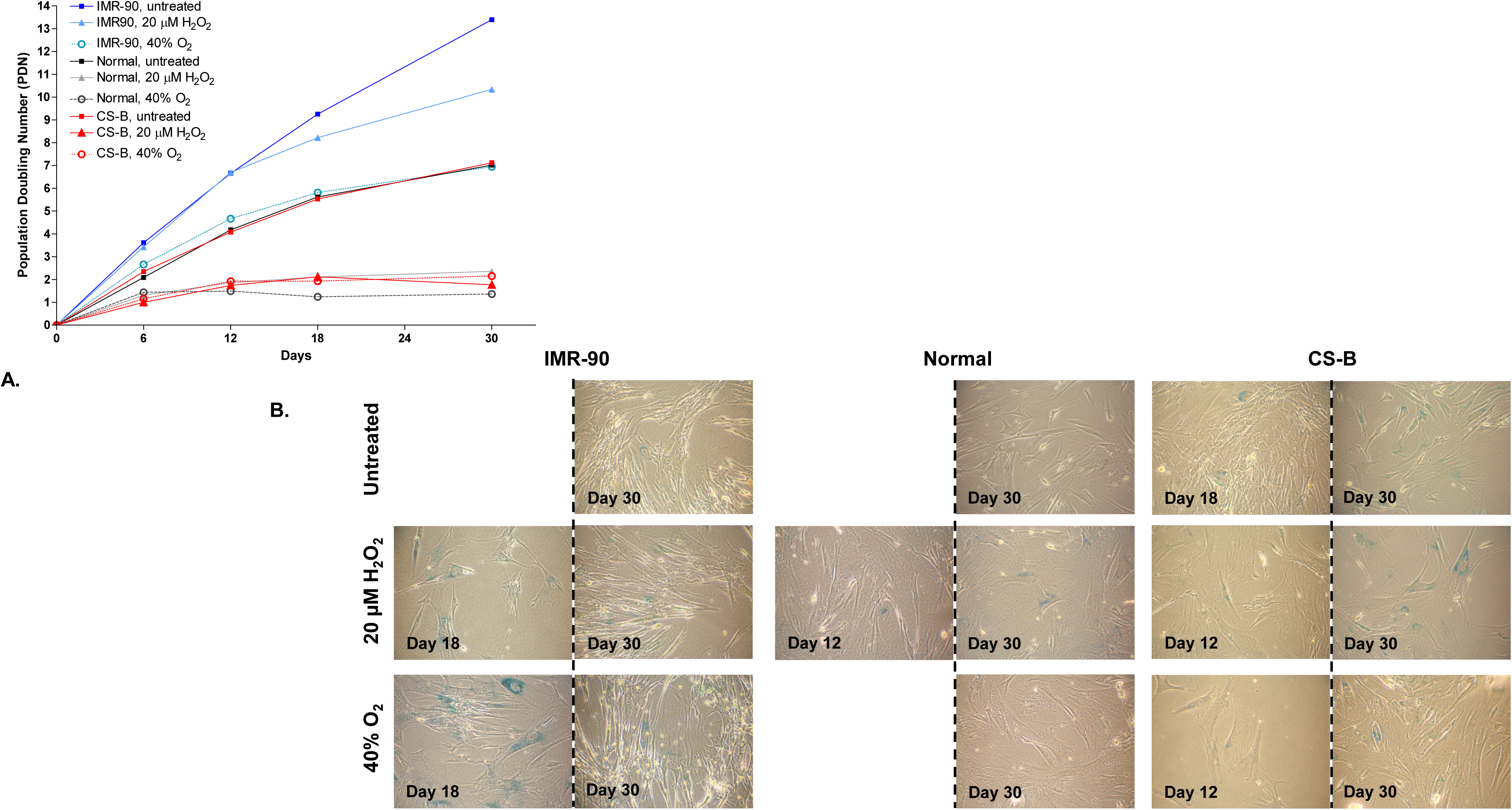
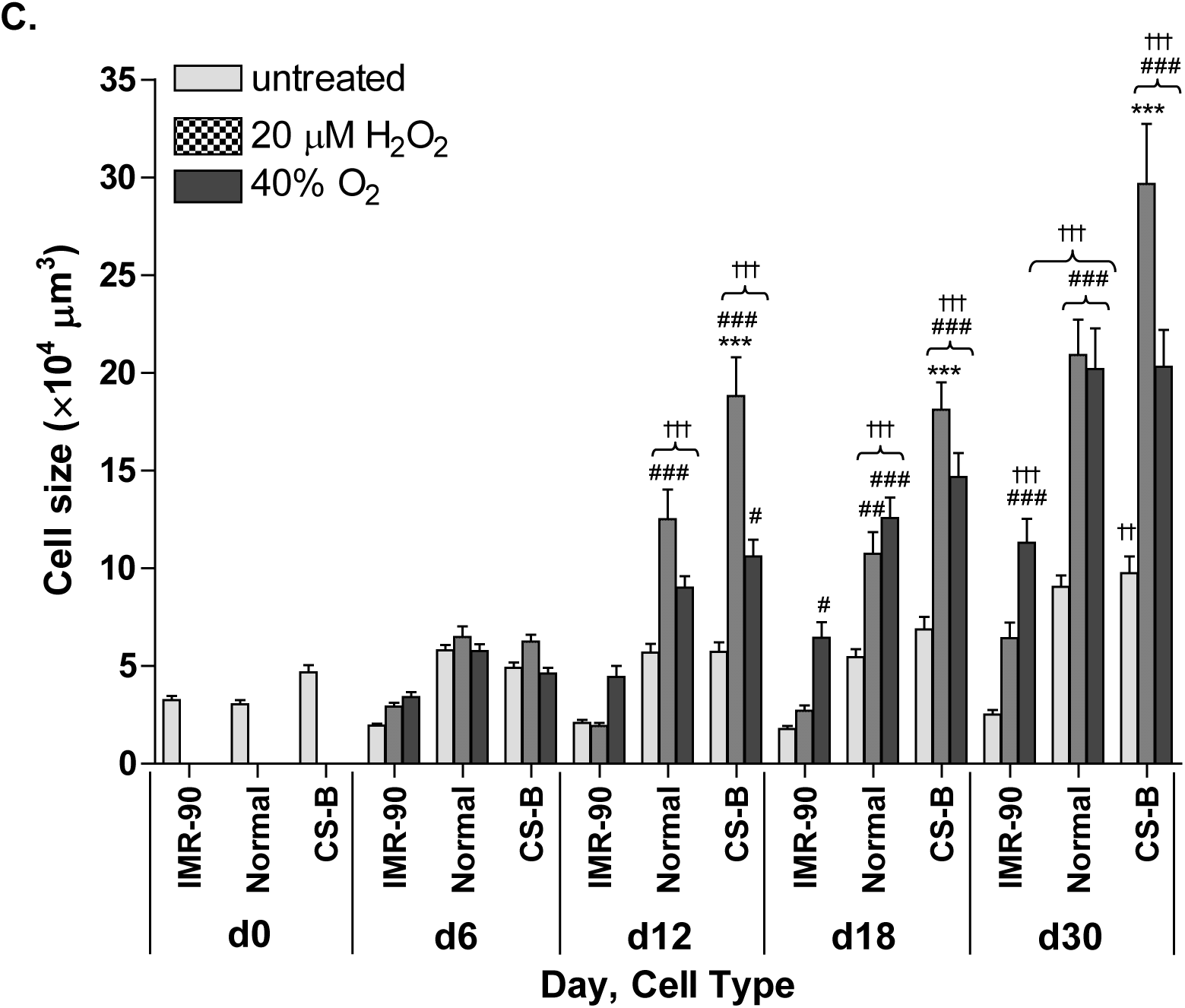
**A)** Growth kinetics of fibroblasts as depicted by population doubling number (PDN) under chronic oxidative stress of 30 days. PDN is determined by cell counting at intervals when untreated cells reach 90% confluence. Cells under oxidative stress show reduced PDN compared to that of untreated counterparts. CS-B and *Normal* cells display similar growth kinetics with or without oxidative stress. **B)** Expression of senescence-associated beta-galactosidase (SA ß-gal) at 100× magnification, as depicted by blue stains Shown are pictures of first day of detectable stains and last day of treatment (Day 30). Where the stain is undetectable before the end of treatment, no picture is shown. Positive staining is detected earlier in cells under oxidative stress when compared to that of cells grown under standard conditions. CS-B cells grown in 40% O_2_ exhibit positive staining earlier than control counterparts. **C)** Cell volume (× 104 µm3) under chronic oxidative stress at specified days. ***P < 0.001 compared to *Normal* cells; #P < 0.05, ##P < 0.01, ###P < 0.001 compared to the untreated counterparts, ††P < 0.001, †††P < 0.001 compared to the size of that of the start of the treatment (day 0). Data is represented as mean ± S.E. Two-way ANOVA was used in the statistical analysis.

Senescence was further investigated using other parameters such as changes in cell morphology indicative of senescence, expression of senescence-associated beta-galactosidase (SA β-gal), as well as increase in cell volume. Both *Normal* and CS-B cells grown under standard culture conditions started to exhibit morphological changes reminiscent of senescence by day 27 data not shown) while IMR-90 cells did not exhibit senescent morphology throughout the experimental period. Chronic exposure to 20 µM H₂O₂ and 40 % O_2_ both hastened the onset of senescent morphological features of all cell types, with CS-B cells showing these features earlier than both IMR-90 and *Normal* fibroblasts. Specifically, following 20 µM H_2_O_2_ treatment, IMR-90, *Normal* and CS-B cells showed characteristics of senescent morphology on days 17, 12 and 7, while these cells, when grown in 40 % O2, exhibited similar changes on days 11, 17 and 7, respectively. Cells were also stained for SA ß-gal depicted by blue stain during the process of the 30-day treatment (**Figure 4B**).CS-B fibroblasts exhibited positive staining as early as the 18th day of treatment (**Figure 4B**), while such positive staining was not detected even after 30 day s in *Normal* fibroblasts and minimal positive staining for IMR90 cells at 30 days of culturing. Blue staining was accelerated in all three cell types when they were chronically exposed to oxidative stress. When exposed chronically to 20 µM H_2_O_2_, IMR-90 fibroblasts exhibited positive staining on day 18, while *Normal* and CS-B fibroblasts stained positive earlier, on day 12. Interestingly, *Normal* cells exposed to 40% O_2_ did not stain positive even on day 30 (**Figure 4B**) despite showing morphology indicative of senescence, such as cell enlargement, flattening and granulation (data not shown). Under this condition, IMR-90 cells stained positive for SA ß-gal on day 18, but CS-B cells showed positive staining earlier, at day 12 (**Figure 4B**).

Six days into exposure to chronic oxidative stress, cell volumes had not increased significantly, nor were there significant differences between cell types or treatments (**Figure 4C**). Untreated IMR-90 fibroblasts did not exhibit any changes in size with time, but both untreated *Normal* and CS-B cells on the last day of treatment were larger than day 0 (P < 0.001). IMR-90 cells exposed chronically to 20 µM H_2_O_2_ did not exhibit significant changes in cell volumes with time. However, both *Normal* and CS-B fibroblasts treated with 20 µM H_2_O_2_ showed a volume increase when compared to their untreated (P < 0.01) and day 0 (P< 0.001) counterparts from day 12. Notably, under this treatment condition, CS-B cells from day 12 onwards were significantly larger in volume than that of *Normal* cells (P < 0.001). Chronic exposure to 40 % O_2_ resulted in a significant increase in cell volume in *Normal* (P < 0.001) and IMR-90 (P < 0.05) fibroblasts at day 18 and CS-B cells at day 12 (P < 0.05) compared to their untreated counterparts. Also, under this condition, a significant increase in cell volume compared to day 0 was seen in IMR-90 fibroblasts at day 30, *Normal* and CS-B cells at day 12 (P < 0.001).

We next looked at TRF lengths of *Normal* and CS-B cells in the beginning and end of the treatment. An image of the resulting southern blot is shown (**Figure 5A**). *Normal* and CS-B cells showed a decrease in TRF length regardless of condition after 30 days (**Figure 5B**). Importantly, both cell types when exposed to oxidative stress of either 20 µM H_2_O_2_ and 40 % O_2_ saw an increased loss of TRF length compared to that of their counterparts grown under standard conditions (**Figure 5C**). To note, CS-B cells had longer telomeres than that of *Normal* cells and showed an augmented reduction in TRF lengths than *Normal* cells regardless of condition type. Taking PDN into account, we analysed the rate of TRF attrition and found that the rate of attrition of TRF of cells under oxidative stress was greatly increased compared to that of untreated controls (**Figure 5C**). Interestingly, CS-B cells exhibited spontaneous telomere shortening as seen by the increased TRF loss in untreated cells compared to *Normal* cells. While CS-B cells, regardless of treatment condition, displayed an increased rate of TRF attrition compared to *Normal* cells, those treated with 20 µM H_2_O_2_ showed a stark difference to that of *Normal* cells (**Figure 5C**).

**Figure 5:**
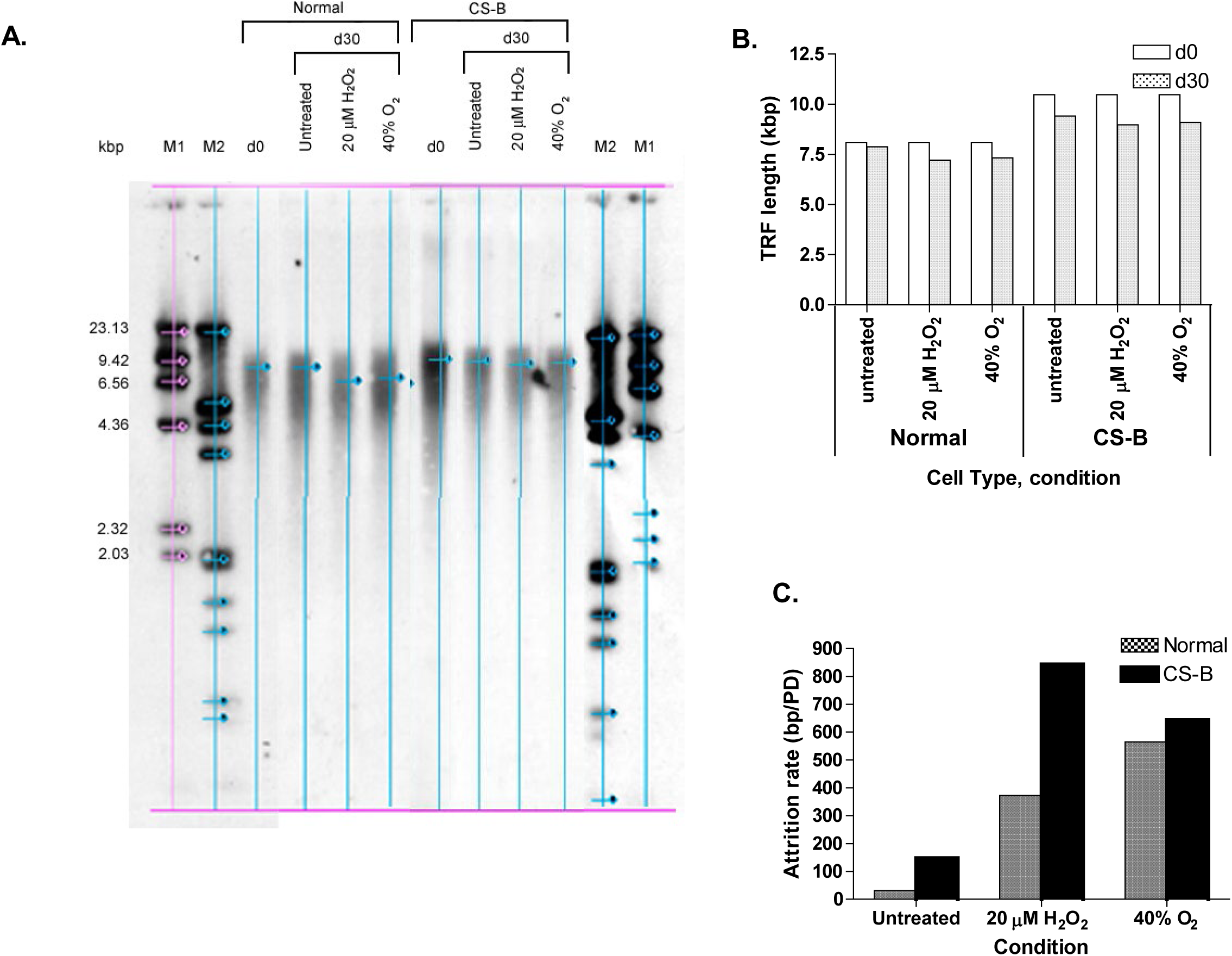
**A)** Southern blotting of TRF obtained from a genomic DNA digest using Hinf I and Rsa I restriction enzymes. **B)** TRF length at start of treatment and following 30 days under chronic oxidative stress. TRF length decreases for both *Normal* and CS-B cells under all conditions following 30 days. **C)** Rate of TRF length attrition following 30 days of chronic oxidative stress. TRF attrition rate is derived by dividing the decrease of TRF (**B**) by PDN (**5A**). Attrition rate increases in both *Normal* and CS-B cells following chronic oxidative stress. CS-B cells displayed an increased attrition rate compared to normal counterparts.

### 3.7 Differential gene expression patterns in CSB-deficient and *Normal* fibroblasts following H_2_O_2_ treatment

Microarray was performed on fibroblasts exposed to 40 µM H_2_O_2_ 2 h after exposure and fibroblasts that were allowed to recover after the 2 h in fresh medium without H_2_O_2_ for 22 h. The 2-D hierarchical clustering of the analysis of the treated cells versus the untreated cells at the two time points revealed differential gene expression patterns between CSB-deficient and *Normal* fibroblasts following H_2_O_2_ treatment (**Figure 6A**). Gene ontology interrogation has separated these differentially expressed genes into cellular processes (**Figure 6B).** Genes representing three main clusters: cell death, DNA damage repair and cell cycle regulation are listed in **Table 2**.

**Figure 6.**
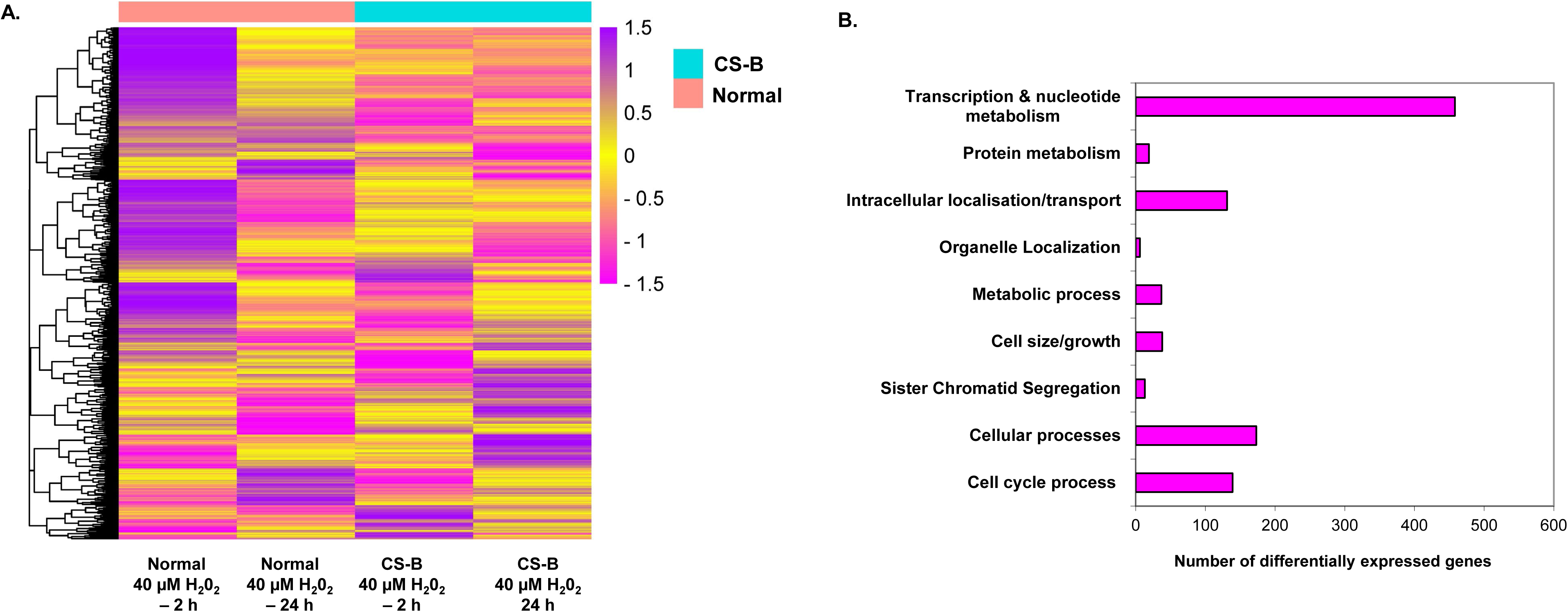
Microarray of fibroblasts treated with 40 µM H_2_O_2_ for 24 h. **A.** Hierarchical Cluster displaying gene expression changes. Data is expressed as treated cells compared to untreated control counterparts. N depicts normal fibroblasts; C depicts CS-B fibroblasts. **B.** Brief overview of the number of genes differentially expressed in categorised biological processes of interest following H_2_O_2_ treatment. A total of 1047 genes were differentially expressed.

**Table 2.** Functional groupings/clusters of differentially expressed genes in fibroblasts immediately following H_2_O_2_ treatment (2h) and 22h after recovery (24 h).

| Gene Name | Genbank ID | Normal 2h | Normal 24 h | CS-B 2h | CS-B 24h |
| --- | --- | --- | --- | --- | --- |
| **Cell death** |  |  |  |  |  |
| *TRAF7* | NM_032271 | -1.02 | -1.16 | **1.54** | 1.03 |
| *DDIT3* | NM_004083 | **3.86** | **2.01** | **1.95** | 1.36 |
| *TP53I3* | NM_147184 | -1.04 | **2.19** | -1.01 | 1.22 |
| *TP53BP2* | NM_005426 | -1.35 | -1.17 | <u>-1.56</u> | -1.06 |
| *SPHK1* | NM_021972 | 1.37 | <u>-2.73</u> | 1.05 | 1.19 |
| *Survivin/API4* | NM_001168 | -1.26 | -1.22 | 1.05 | <u>-3.49</u> |
| *CDKN1A/p21Cip1* | NM_078467 | 1.18 | **2.07** | 1.31 | 1.44 |
| **DNA damage repair** |  |  |  |  |  |
| *POLE* | NM_006231 | -1.08 | -1.12 | **1.81** | -1.41 |
| *POLE2* | NM_002692 | -1.25 | <u>-1.82</u> | -1.01 | <u>-1.74</u> |
| *RFC3* | NM_002915 | <u>-1.50</u> | -1.07 | -1.06 | <u>-1.83</u> |
| *RFC4* | NM_002916 | -1.18 | <u>-1.81</u> | 1.07 | <u>-1.76</u> |
| *RFC5* | NM_007370 | -1.06 | -1.19 | -1.00 | <u>-1.70</u> |
| *LIG1* | NM_000234 | -1.18 | 1.23 | 1.31 | <u>-1.73</u> |
| *XPC* | NM_004628 | -1.15 | **1.84** | **1.55** | 1.14 |
| *EXO1* | NM_003686 | <u>-1.57</u> | <u>-1.58</u> | 1.17 | <u>-1.99</u> |
| *RAD51* | NM_002875 | -1.14 | <u>-1.66</u> | -1.13 | <u>-1.75</u> |
| *RAD54L* | NM_003579 | 1.02 | -1.03 | 1.15 | <u>-1.52</u> |
| *GSTE1* | NM_016426 | -1.35 | -1.01 | 1.23 | <u>-2.34</u> |
| *H2AFX* | NM_002105 | 1.27 | -1.08 | 1.44 | <u>-2.46</u> |
| *BLM* | NM_000057 | -1.24 | -1.25 | -1.01 | <u>-1.54</u> |
| *GADD45B* | NM_015675 | **2.49** | <u>-1.61</u> | **1.65** | -1.23 |
| *GADD45G* | NM_006705 | **1.63** | 1.01 | 1.23 | 1.07 |
| **Cell cycle regulation** |  |  |  |  |  |
| *GADD45B* | NM_015675 | **2.49** | <u>-1.61</u> | **1.65** | -1.23 |
| *GADD45G* | NM_006705 | **1.63** | 1.01 | 1.23 | 1.07 |
| *CDT1* | NM_030928 | -1.01 | <u>-2.74</u> | 1.44 | <u>-2.05</u> |
| *CDC45L* | NM_003504 | -1.37 | <u>-2.04</u> | 1.17 | <u>-2.21</u> |
| *POLE* | NM_006231 | -1.08 | -1.12 | **1.81** | -1.41 |
| *RBL1* | NM_002895 | -1.34 | -1.14 | 1.10 | <u>-1.76</u> |
| *E2F2* | NM_004091 | -1.13 | -1.44 | 1.21 | <u>-1.89</u> |
| *p19* | NM_001800 | 1.21 | 1.01 | 1.02 | -1.36 |
| *CDKN1A/p21Cip1* | NM_078467 | 1.18 | **2.07** | 1.31 | 1.44 |
| *CDKN3* | NM_005192 | -1.10 | 1.48 | -1.13 | <u>-3.68</u> |
| *CDK2* | NM_001798 | -1.18 | -1.13 | 1.03 | <u>-1.67</u> |
| *Cyclin E2* | NM_057735 | -1.05 | <u>-1.80</u> | 1.20 | <u>-1.74</u> |
| *CDC2* | NM_001786 | -1.02 | 1.36 | 1.03 | <u>-2.30</u> |
| *Cyclin A2* | NM_001237 | -1.32 | -1.09 | 1.01 | <u>-3.58</u> |
| *Cyclin B1* | NM_031966 | -1.16 | -1.29 | 1.00 | <u>-2.33</u> |
| *Cyclin B2* | NM_004701 | -1.19 | **1.51** | -1.05 | <u>-3.30</u> |
| *CDC25A* | NM_001789 | 1.08 | <u>-2.40</u> | 1.14 | -1.49 |
| *CDC25B* | NM_004358 | -1.11 | 1.10 | 1.16 | <u>-1.99</u> |
| *CDC25C* | NM_001790 | -1.04 | 1.17 | 1.08 | <u>-2.02</u> |
| *WEE1* | NM_003390 | 1.23 | -1.45 | 1.26 | <u>-1.60</u> |
| *CDC20* | NM_001255 | -1.22 | -1.48 | -1.06 | <u>-4.22</u> |
| *KNTC1* | NM_014708 | -1.15 | 1.01 | 1.07 | <u>-1.84</u> |
| *CDCA8* | NM_018101 | -1.15 | -1.15 | 1.26 | <u>-3.61</u> |
| *BUB1* | NM_004336 | -1.09 | -1.11 | 1.21 | <u>-3.69</u> |
| *BUB1B* | NM_001211 | -1.21 | -1.04 | -1.05 | <u>-1.53</u> |
| *KIF20B* | NM_016195 | -1.34 | -1.02 | -1.05 | <u>-2.96</u> |
| *PRC1* | NM_003981 | -1.35 | 1.43 | 1.19 | <u>-3.28</u> |
| *AURKA* | NM_198434 | -1.05 | -1.03 | 1.11 | <u>-3.63</u> |
| *CDC14A* | NM_033313 | <u>-1.51</u> | 1.16 | -1.31 | -1.14 |
| *UBE2C* | NM_181799 | 1.01 | -1.01 | -1.02 | <u>-1.69</u> |

## 4.0 Discussion

CS-B primary fibroblasts in our study have shown to be less susceptible to H_2_O_2_-induced cell death compared to control cells, possibly because these cells have a functional defect in the TCR pathway. Moreover, aberrant repair pathways may result in signalling changes that incapacitate damage-induced cell death [7–9, 30, 31]. Contrary to our findings that CSB-deficient cells appeared less sensitive to oxidative stress, other groups have found CSB-deficient cells to be hypersensitive to UV-induced apoptosis [32–35]. However, CSB-deficient mouse embryonic fibroblasts and embryonic stem cells have shown differential sensitivity to UV and ionizing radiation [33] and the sensitivity of CS cells to oxidative stress has been suggested to be an inconsistent feature [10]. Thus, it is possible that CS-B human fibroblasts might respond with differing sensitivity to H2O2 compared to UV. In support of this, CSB-deficient mouse keratinocytes have presented with increased resistance to UVB-induced apoptosis accompanied by prolonged growth arrest compared to NER-deficient keratinocytes [36]. Given more time however, we may see cell death or more prominent cell cycle arrest in the CS-B fibroblasts… ROS has been shown to induce DNA damage resulting in obligatory cell cycle arrest [37, 38]. Our data on increased sub-G1 (data not shown) and decrease in cell viability following H_2_O_2_ exposure supports these earlier concepts and observations.

Morphology of the fibroblasts 2 h immediately following H_2_O_2_ treatment revealed that *Normal* cells are indeed more sensitive to H_2_O_2_ as shown by the refractive cells. The cell cycle patterns pertaining to sub-G1 population support our cell viability data. Although CS-B cells neither displayed much cell death nor significant changes in the cell cycle, they exhibited a slight late S phase/G2/M phase arrest with time. Such observation may be either attributed to insufficient damaged induced by the used concentrations of H_2_O_2_ or to defective cell cycle checkpoint activation in CS-B cells. While the latter is a very probable reason, loss of GG-NER but not TCR has been found to affect checkpoint triggering [39]. Cells exposed to oxidative stress in culture accumulate protein modifications, lipid peroxidation and DNA damages in the form of strand breaks and oxidized bases [40–43]. [41, 44–46]. Accumulation of oxidative DNA lesions and reduced efficiency of DNA repair has been associated with ageing phenotypes and cancer [47, 48]. Our observation on higher presence of micronuclei and increased/persisted DNA damage (comet assay) indicates the role of CSB in the repair of oxidative lesions. In line with this, organs of CSB-deficient mice and brains of CS patients have been found to harbour enhanced oxidative DNA lesions [49–52]. Treatment with low levels of H_2_O_2_ sufficed to induce significantly more DNA damage in CS-B fibroblasts than in *Normal* fibroblasts, supporting the notion that a compromised TCR renders CS-B cells prone to more severe consequences on the genome when faced with external stresses. Not only are CS-B cells more susceptible to H_2_O_2_-induced DNA damage as depicted by augmented tail moment lengths at the 2-h time point, but they are also deficient in competent repair of the incurred damage as depicted by the lack of reduction of tail moments back to that of basal levels. further corroborating that these cells accumulate oxidative stress and that CSB and the TCR are involved in the eradication of oxidative DNA lesions. To note, although CSB has been established to be involved in oxidative stress repair and BER, which is the characterised repair pathway for dealing with oxidative damage [53–56], It is not surprising that both *Normal* and CS-B fibroblasts are able to reverse tail moment lengths by a considerable extent during recovery.

Our data demonstrates that deficiency in CSB that incapacitates TCR leads to cell cycle arrest and increased vulnerability to oxidative stress and accumulation of oxidative damage. This is expected, with the availability of evidence pointing to the involvement of CSB in not only TCR and BER repair pathways, but also recovery from post-UV transcriptional arrest [9, 13, 57–59]. Thus our data corroborates with the currently accepted theory that oxidative stress is a major contributing factor in CS symptoms that cannot be attributed to UV exposure such as neurological disorders, developmental delays and progeria [11, 13, 60, 61].

Oxidative stress has been shown to cause stress-induced premature senescence (SIPS) that possibly drives ageing via induction of stochastic damage, acceleration of telomere attrition and inhibition of proliferation [47, 62–65]. As terminal chromosomal structures, telomeres protect against aberrant recombination and exonuclease attacks on the DNA [66, 67]. Due to the end replication problem, critically short are thought to signal permanent exit of the cell cycle and entry into senescence via a DNA damage response [21, 68, 69]. Telomeres exhibit increased susceptibility to single-stranded breaks induced by oxidative stress [70–73]., oxidative stress induces accelerated telomere shortening contributing to SIPS in turn cells, express SA β-gal and cease proliferating.

To mimic physiological chronic oxidative stress, we treated the cells with low dose oxidative stress over 30 days while observing markers of senescence such as PDN, senescent morphology, SA β-gal expression, cell volume and telomere attrition rate. Two experimental set ups were performed, where cells were treated with 10 µM H2O2 for the first set up, and with either 20 µM H2O2 or 40% O2 for the second. PDN kinetics and time of appearance of senescent morphology in untreated CS-B fibroblasts with respect to untreated *Normal* cells were not consistent on the two different occasions. Despite the inconclusiveness of whether loss of CSB in fibroblasts leads to reduced growth capacity or hastened senescent characteristics, untreated CS-B fibroblasts exhibited a significant increase in cell volume after 30 days, a phenomenon that control cells did not show. Also, they stained positive for SA β-gal earlier than control cells and demonstrated spontaneous telomere attrition, indicating that lack of CSB may put cells at risk for premature senescence.

Chronic exposure to 10 µM H_2_O_2_ decreased the PDN slightly in both *Normal* and CS-B fibroblasts, indicating that oxidative damage can slow down proliferative capacity and resulted in hastened onset of senescent morphology in both cell types. Importantly, although the starting length of telomeres in CS-B fibroblasts was longer than that of *Normal* fibroblasts, it was rapidly lost over time regardless of exposure to 10 µM H_2_O_2_, indicating spontaneous telomere attrition. Despite the increased rate of telomere loss in CS-B fibroblasts compared to *Normal* fibroblasts, CS-B cells did not show overt cessation of proliferation and senescent morphology, possibly because the absolute length of the telomeres dictate onset of senescent characteristics such as permanent cell cycle arrest, and these telomeres may not have reached that critical length yet. Treatment with 10 µM H_2_O_2_demonstrated a slight increase in the rate of telomere loss in *Normal* cells compared to that of untreated, consistent with findings that oxidative stress enhances rate of telomere shortening [63–65]. Such a treatment condition also increased the rate of telomere loss in CS-B cells especially in the first 12 days of the treatment, demonstrating that the lack of intact CSB increases the rate of oxidative stress-induced telomere shortening. To note, both untreated and treated CS-B fibroblasts exhibited a largely increased rate of telomere attrition compared to their *Normal* fibroblast counterparts, indicating that the loss of CSB function renders cells more susceptible to telomere loss regardless of oxidative stress, thus implying a possible role of CSB in telomere maintenance.

When cells were exposed to either 20 µM H_2_O_2_ or 40% O_2_, PDN dropped drastically, senescent-like morphology and positive staining of SA ß-gal appeared earlier, cell volume increased earlier, and telomere attrition rates increased drastically compared to untreated counterparts. Specifically, both *Normal* and CS-B fibroblasts exhibited a severe reduction in PDN following chronic exposure to either form of oxidative stress indicative of a halt in the cell cycle. These observations indicate that exposure to oxidative stress chronically can indeed induce SIPS, consistent with previous findings [63–65].

CSB may also be involved in telomere protection akin to the essential role of DNA repair factors such as the NHEJ components, ATM, BLM, DNA-PKcs, WRN and even the NER factor XPF, in the maintenance of telomeric integrity. The function of these proteins as caretakers of both the genome and telomeres implicates them in carcinogenesis, ageing and premature ageing syndromes [22, 23, 74, 75]. A classic example is that of ataxia telangiectasia cells which exhibit accelerated telomere loss [76] and premature senescence [77]. That CS-B fibroblasts in our study exhibited the same features strongly correlates the role of the CSB protein to telomere maintenance and cellular age since the loss of this protein renders cells more sensitive to oxidative SIPS and telomere attrition. Significantly, our findings also implicate oxidative stress as an additional factor in the progeroid phenotypes of CS patients, which corroborates with current knowledge that CSB is not only involved in the TCR of oxidative DNA lesion repair [54], but also in the repair of spontaneous oxidative lesions in non-transcribed genes [78] and BER [8]. Further studies are required to understand the mechanistic role of CSB in telomere regulation.

Microarray was used to ascertain the gene expression profiles following treatment with a sub-lethal concentration of 40 µM H_2_O_2_. In relevance to this work, we looked at select genes in three broad clusters: cell death, DNA damage-repair and cell cycle regulation (Table 2). Here, we have shown that exposure to H_2_O_2_ in CSB-deficient cells resulted in the significant down-regulation of DNA damage repair genes including DNA polymerase complex genes *POLE, POLE2, RFC3, RFC4* and *RFC5*, BER pathway gene *LIG1*, GG-NER-specific gene *XPC* and homologous repair genes *EXO1*, *RAD51* and *RAD54L*. *BLM* helicase and *H2AFX* were also down-regulated in CS-B cells. This is suggestive that the lack of a single functional repair protein in a particular repair pathway can interfere with the repair capacity of other pathways possibly because of crosstalk between repair pathways. As H_2_O_2_ does not only inflict TCR or BER-type lesions, the mis-regulation of genes in other repair pathways could impair the rectification of other lesion types, which is apparent in our damage marker assays where CS-B fibroblasts exhibited significantly more damage than *Normal* cells. Interestingly, , the loss of CSB resulting in the down-regulation of *BLM* is of interest as *BLM* helicase has been found to play a role in telomere maintenance [79]. Despite the perceived lack of DNA repair in CS-B cells, these cells were capable of up-regulation of GADD genes at the 2-h time point. Notably, genes associated with the polymerase complex were also downregulated in *Normal* cells at the recovery time point, suggesting that H_2_O_2_ treatment disrupts DNA replication associated with repair and physiological DNA replication regardless of the presence of functional CSB.

Following 22 h recovery from H_2_O_2_ in fresh media, both *Normal* and CS-B fibroblasts exhibited up-regulation of pro-apoptotic genes *DDIT3* and p53-dependent pro-apoptotic gene *TP53I3*. In addition, the anti-apoptotic gene *SPHK1* was down-regulated in *Normal* cells and the anti-apoptotic gene *Survivin* was down-regulated to a larger extent in CS-B cells than *Normal* cells. This indicates that both *Normal* and CS-B fibroblasts are predisposed to cell death when treated with 40 µM H_2_O_2_.Incidentally, such differential expression supports the cell viability in *Normal* cells (Figure 1). Notably, apoptosis is a complex process and to ascertain why CS-B cells did not exhibit cell death despite these expression patterns requires more investigation that the scope of this study can provide. Also, as numerous studies have demonstrated the susceptibility of CSB-deficient cells to oxidative stress-induced cell death, it is possible that these cells may require a longer time before they exhibit cell death.

Microarray reflected that many cycle genes were downregulated 22 h after recovery from H2O2 treatment. The downregulation of pro-replicative genes *CDT1, CDC45L*, *CDC14A* and *POLE*, and *E2F2* which is required for gene transcription for S phase entry indicates that 40 µM H2O2 is sufficient to interfere with DNA replication. Cyclins, cdks such as *CDK2* and *CDC2*, CDC25 phosphatases required to activate cdks, ubiquitin-associated genes *CDC20* and *UBE2C*, and mitotic genes such as *KNTC1*, *CDCA8*, *BUB1*, *BUB1B*, *KIF20B*, *PRC1* and *AURKA* were also significantly down-regulated. These, together with the up-regulation of the negative regulator cdk inhibitor, *CDKN1A/p21^Cip1^*, in CS-B cells indicate that the lack of CSB results in cell cycle arrest after H2O2 treatment. In support of our gene expression study, CSB is known to be involved in transcription. Not surprisingly, defects in this gene resulting in its loss of function have been shown to alter gene expressions. Some of these genes found were involved in DNA repair and cell cycle regulation [1, 14].

In this study, we have shown that the loss of CSB increases susceptibility to H_2_O_2_-induced genotoxicity and compromises repair capacity. Consistent with previous studies, we have also demonstrated that chronic oxidative stress accelerates premature senescence characteristics. In addition, CS-B fibroblasts exhibit premature senescent features, whose manifestations were accelerated upon exposure to chronic oxidative stress. Here, we provide evidence that the symptoms of CS, such as premature ageing resulting from the loss of regenerative capacity of organs and developmental delays, may not only be due to apoptosis from transcription block or damage but very possibly from SIPS resulting from endogenous stress that leads to telomere loss and stochastic damage. Interestingly, cells defective in CSB have demonstrated increased breaks in their DNA even when grown in ambient oxygen [1], suggesting that increasing their oxidative load may further augment the number of breaks. These breaks may well be at the telomeres, hence potentiating spontaneous telomere shortening, as seen in this study. Being a protein that bears the burden of multiple functions such as the TCR, BER, transcription and chromatin remodelling [8]., it is no surprise that the loss of CSB function renders cells incapable of repair and transcription, which ultimately incapacitates the normal functioning of the cell, leading to cell death or senescence. Thus, our study implicates the role of CSB in the repair of oxidative stress and maintenance of telomeres.

## 5.0 Conclusion

This study elucidates the complex role of Cockayne Syndrome B (CSB) protein in cellular responses to oxidative stress, highlighting its dual impact on maintaining cell viability under acute conditions and mediating susceptibility to chronic oxidative environments. Our findings demonstrate that CSB-deficient fibroblasts exhibit an accelerated rate of telomere attrition and premature senescence under prolonged oxidative stress, underscoring the significance of CSB in genome maintenance and aging processes relevant to the pathology of Cockayne Syndrome. While the robust in vitro evidence from fibroblast models provides valuable insights, the cellular specificity of CS pathology and the dynamic in vivo environment may influence these outcomes, suggesting broader investigative approaches are needed. Future research should explore the functional dynamics of CSB in varied cell types, particularly neuronal cells, and extend these studies into in vivo models to validate therapeutic targets. Further elucidation of CSB’s interactions within other DNA repair pathways could also enhance our understanding of its broader role in genomic integrity and inform the development of interventions for CS and related premature aging disorders.

## Acknowledgements

MPH acknowledges the assistance from Genome Stability Laboratory members during this study.

